# Caveola mechanotransduction reinforces the cortical cytoskeleton to promote epithelial resilience

**DOI:** 10.1101/2023.03.29.534729

**Authors:** John W. Brooks, Vikas Tillu, Suzie Verma, Brett M. Collins, Robert G. Parton, Alpha S. Yap

## Abstract

As physical barriers, epithelia must preserve their integrity when challenged by mechanical stresses. Cell-cell junctions linked to the cortical cytoskeleton play key roles in this process, often with mechanotransduction mechanisms that reinforce tissues. Caveolae are mechanosensitive organelles that buffer tension via disassembly. Loss of caveolae, through caveolin-1 or cavin1 depletion, causes activation of PtdIns(4, 5)P_2_ signalling, recruitment of FMNL2 formin, and enhanced cortical actin assembly. How this equates to physiological responses in epithelial cells containing endogenous caveolae is unknown. Here we examined the effect of mechanically-inducing acute disassembly of caveolae in epithelia. We show that perturbation of caveolae, through direct mechanical stress, reinforces the actin cortex at adherens junctions. Increasing interactions with membrane lipids by introducing multiple phosphatidylserine-binding undecad cavin1 (UC1) repeat domains into cavin1 rendered caveolae more stable to mechanical stimuli. This molecular stabilization blocked cortical reinforcement in response to mechanical stress. Cortical reinforcement elicited by the mechanically-induced disassembly of caveolae increased epithelial resilience against tensile stresses. These findings identify the actin cortex as a target of caveola mechanotransduction that contributes to epithelial integrity.

## Introduction

Epithelia form many of the biological barriers of the metazoan body. They mediate regulated secretion and absorption and also protect the body from its external environment. Central to this physiological function is the integrity of cell-cell cohesion, which is mediated by multiple specialised intercellular junctions, where cell-cell adhesion systems are linked to elements of the cytoskeleton. For example, in adherens junctions (AJ), the E-cadherin complex physically and functionally engages with the actomyosin cytoskeleton (Charras & Yap, 2018; Mège & Ishiyama, 2017), whereas in desmosomes, desmogleins and desmocollins associate with the intermediate filament cytoskeleton (Delva *et al*, 2009).

Ultimately, these adhesion systems must preserve epithelial integrity by resisting mechanical forces that can disrupt cell-cell cohesion. These disruptive forces can be applied from outside the tissue, as is associated with physical trauma or pulmonary inflation (Zhong *et al*, 2020), and also arise from within the tissue, when the actomyosin cytoskeletons of epithelial cells contract against the AJ that connect cells together (Charras & Yap, 2018). To cope with these challenges, epithelial tissues have evolved a variety of mechanisms to both sense and respond to mechanical forces (Choi *et al*, 2012; Haas *et al*, 2020; Huveneers *et al*, 2012; Pokutta *et al*, 2002; Spadaro *et al*, 2017; Yao *et al*, 2014; Yonemura *et al*, 2010). For example, the cadherin molecular complex in AJ contains mechanosensitive elements, such as the adhesive binding domain of E-cadherin, the cytosolic adapter protein, α-catenin, and the cadherin-associated protein, Myosin VI. Mechanical forces transmitted via E-cadherin can induce conformational changes in α-catenin to promote vinculin and F-actin binding (Noordstra *et al*, 2023), while recruitment of Myosin VI engages a signal transduction apparatus that activates RhoA at AJ (Acharya *et al*, 2018).

Caveolae constitute another important means of mechanoprotection in many tissues, being particularly abundant in tissues which experience significant mechanical forces. Caveolae consist of plasma membrane invaginations that are created when caveolin (Cav) integral membrane proteins engage with cytosolic cavin coat proteins. Caveolae are distributed at sites of cell-cell contact (Lu *et al*, 2003; Orlichenko *et al*, 2009; Palacios *et al*, 2002; Volontè *et al*, 1999) and are clustered at AJs in epithelial cells (Teo *et al*, 2020). Caveola mechanosensing is currently understood to be mediated by the disassembly of cavins, leading to the partial or complete flattening of caveolae (Sinha *et al*, 2011). Caveola-deficient animals are prone to injury due to mechanical stress, as has been observed in the notochord of cavin 1b-deficient zebrafish embryos (Garcia *et al*, 2017; Lim *et al*, 2017), in endothelia (Cheng *et al*, 2015) and in skeletal muscle (Lo *et al*, 2016). The best understood mechanism for mechanoprotection involves the release of membrane when caveolae disassemble, which serves to buffer against increases in membrane tension (Cheng *et al*., 2015; Lo *et al*., 2016; Rog-Zielinska *et al*, 2021).

But caveolae can also interact with the actin cytoskeleton (Echarri & Del Pozo, 2015) and modulate cell signalling to reorganise the cytoskeleton (Radel & Rizzo, 2005; Teo *et al*., 2020) and RhoA-dependent contractility (Goetz *et al*, 2011; Grande-García *et al*, 2007; Hetmanski *et al*, 2019; Yang *et al*, 2011). Previously, we showed that chronic depletion of caveolae, by RNAi directed against either CAV1 or cavin1, causes increased cortical tension in epithelial monolayers through a pathway involving PtdIns(4, 5)P_2_, the formin-like protein FMNL2, and the actin cytoskeleton (Teo *et al*., 2020). However, it was unclear whether this pathway contributed to a physiological role for caveolae when they disassembled in response to mechanical stimulation. Accordingly, in this study we sought to test if cortical regulation is affected when mechanosensing induces the acute disassembly of caveolae, and define the role that this may play in epithelial mechanoprotection. Developing a new molecular strategy to stabilize caveolae, we show that mechanically-induced caveolar disassembly promotes reinforcement of the actomyosin cortex, rapidly altering the mechanical profile of epithelial monolayers to confer resistance to physical insults.

## Results

### Mechanical stresses reinforce the junctional cytoskeleton in a caveola-dependent fashion

To investigate how caveola mechanosensing might affect the cortical cytoskeleton, we applied hypo-osmotic stimulation to confluent MCF-10A mammary epithelial monolayers. Hypo-osmotic stress is a commonly-used manoeuvre that induces the release of cavins from caveola by increasing membrane tension as cells swell (Sinha *et al*., 2011). Monolayers were exposed to growth media with an osmolarity of 150 mOsm L^-1^ (approximately 50% that of physiological conditions) for 5 minutes. Immunofluorescence analysis showed that this reduced junctional staining for cavin1 (relative to the cytoplasmic signal) without affecting Cav-1 localisation, consistent with a disassembly of caveolae (Fig. S1a-e).

Under baseline conditions, control MCF-10A monolayers displayed prominent F-actin (phalloidin) staining at cell-cell junctions, and this was reversibly increased by ∼20% upon short-term application of hypo-osmotic media (Fig. 1a-b and Fig. S2a). This cortical response required caveolae, since Cav-1 shRNA (knock-down, KD) monolayers had increased junctional F-actin at baseline, as previously reported (Teo *et al*., 2020), but this did not increase upon hypo-osmotic stimulation. These differences were specific for Cav-1, as they were restored upon reconstitution of KD cells with exogenous Cav-1 (Fig. 1a-b and Fig. S2a). Myosin IIA was most prominent at the basal regions of control cells prior to hypo-osmotic stimulation and increased at junctions upon stimulation (Fig. 1c). However, this increase in junctional Myosin IIA was not affected by Cav-1 KD, suggesting that it was due to a caveola-independent effect of hypo-osmotic stimulation.

**Figure 1.**
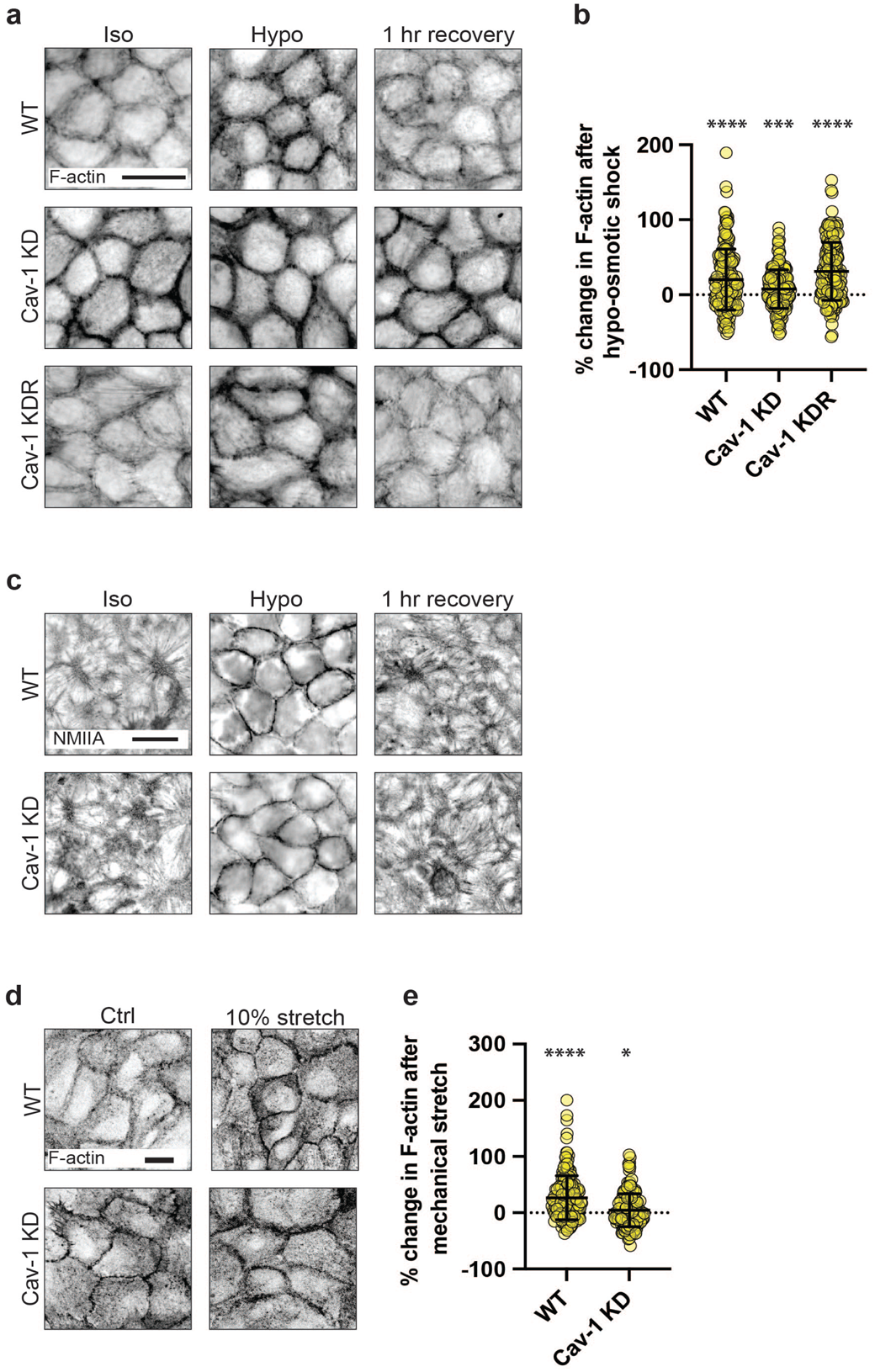
The cytoskeleton at adherens junctions is reinforced by mechanical stimulation of caveolae. **(a)** Fluorescence imaging of F-actin in WT, Cav-1 KD, and Cav-1 KDR MCF-10A monolayers (scale bar = 10 μm). **(b)** Quantification of (a) showing the percentage change in cortical F-actin levels following hypo-osmotic shock. F-actin in WT monolayers was increased by ∼20% (p = <0.0001), Cav-1 KD ∼7% (p = 0.0125), and Cav-1 KDR ∼31% (p = <0.0001). **(c)** Immunofluorescence (IF) imaging of non-muscle myosin II (NMII) type A in WT and Cav-1 KD MCF-10A monolayers after hypo-osmotic shock and following a 1 hr recovery period in iso-osmotic growth media. **(d)** Fluorescence imaging of F-actin in WT and Cav-1 KD MCF-10A monolayers prior to, and following, 10% mechanical stretch (scale bar = 10 μm). **(e)** Quantification of (d) shows that cortical F-actin is increased by ∼27% and ∼4% in WT (p = <0.0001) and Cav-1 KD (p = 0.02238) monolayers following stretch, respectively. All data are means ± SD. All statistical analyses calculated from N=3 independent experiments using unpaired t-tests. Points on graphs represent individual cell junctions. N.s, not significant; *p<0.05; **p<0.01, ***p<0.001; ****p<0.0001.

We next tested if other mechanical stresses that impinge on caveola mechanosensing and disassembly could also elicit cytoskeletal changes. External stretch was applied to confluent MCF-10A monolayers grown on flexible silicon-based substrata in BioFlex® culture plates. Cyclic biaxial mechanical stretch (10% strain, 1 Hz, 5 min) displaced cavin1 from the junctional membranes, suggesting that caveolae were being disassembled (Fig. S1f-h). This was accompanied by a significant increase in junctional F-actin (Fig. 1d-e and Fig. S2b). This cortical response to mechanical stretch was compromised, however, by Cav-1 KD, suggesting a role for caveolae, as we had observed for hypo-osmotic stimulation. Overall, this suggested that caveolae might regulate the junctional cytoskeleton as part of a mechanotransduction pathway.

### Caveola disassembly increases mechanical tension at adherens junctions in response to hypo-osmotic stimulation

We next asked if the increase in junctional F-actin was accompanied by changes in junctional mechanics, focusing on AJ, which are often sites of mechanical tension. Junctional tension was evaluated first by measuring the initial recoil of vertices after laser-ablation of the bicellular AJ that connect those vertices (Liang *et al*, 2016; Michael *et al*, 2016; Ratheesh *et al*, 2012). Control monolayers showed a baseline initial recoil, consistent with there being pre-existing mechanical tension in the bicellular junctions. However, initial recoil was increased significantly after acute hypo-osmotic stimulation and restored upon addition of iso-osmotic media (Fig. 2a). This suggested that hypo-osmotic stimulation could reversibly increase AJ tension. However, hypo-osmotic stress did not increase AJ tension in Cav-1 KD cells (Fig. 2a), although these cells had an elevated baseline tension, as previously reported (Teo *et al*., 2020). This suggested that caveolae might be required for the rapid increase in AJ tension that occurred upon hypo-osmotic stimulation.

**Figure 2:**
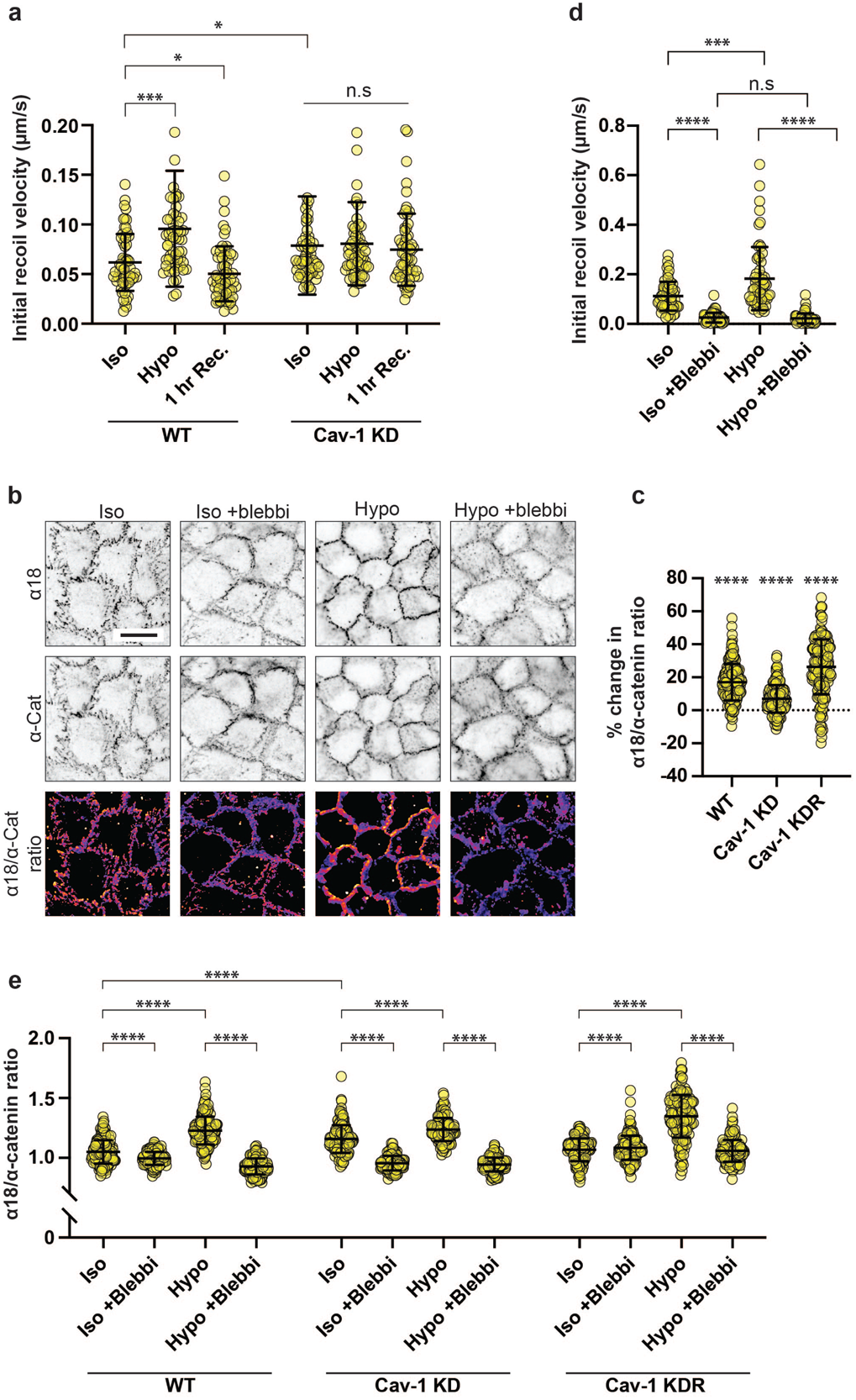
Mechanical activation of caveolae increases tension at adherens junctions. **(a)** Initial recoil velocity (IRV) of cell vertices following junctional laser ablation in confluent MCF-10A epithelial monolayers. In WT monolayers, hypo-osmotic shock increased IRV by ∼55% (p = 0.0006), which is returned to baseline levels following a 1 hr recovery period in iso-osmotic growth medium (p = 0.0306). In monolayers lacking caveolae (Cav-1 KD), baseline tension is increased by ∼28% compared to WT monolayers. Importantly, this is unchanged following hypo-osmotic shock (p = 0.7896). **(b)** IF imaging of total α-catenin (top) and α-catenin in its open conformation (α18) (middle). Lower row shows an image calculation of α18/α-catenin. **(c)** Quantification of (b) showing the percentage changes in the α18/α-catenin ratio following hypo-osmotic shock in WT (∼17%; p = <0.0001), Cav-1 KD (∼7%; p = <0.0001), and Cav-1 KDR (∼26%; p = <0.0001) monolayers. Cav-1 KD monolayers also show a baseline increase of ∼10%. **(d)** In separate experiments, a 1 hr pre-treatment with 25 nM para-aminoblebbistatin (an inhibitor of actomyosin contractility) attenuated the IRV of cell vertices in WT monolayers by ∼77% under control conditions (p = <0.0001). In the absence of para-aminoblebbistatin, IRV was found to increase by ∼63% following hypo-osmotic shock (p = <0.0001), but this was desensitised with the addition of para-aminoblebbistatin (p = 3920). **(e)** Full dataset of α-18/α-catenin ratio (see figure 2b/c) showing data points from individual cell-cell junctions. All data are means ± SD. All statistical analyses calculated from N=3 independent experiments using unpaired t-tests. Points on graphs represent individual cell junctions. N.s, not significant; *p<0.05; **p<0.01, ***p<0.001; ****p<0.0001.

To corroborate the results of recoil experiments, we then assayed changes in the conformation of the E-cadherin-associated protein, α-catenin, as an index of molecular-level tension at AJ. Application of mechanical tension causes conformational unfolding of α-catenin, exposing cryptic epitopes including one in the central M-domain that is recognised by the α18 monoclonal antibody (mAb) (Noordstra *et al*., 2023; Yonemura *et al*., 2010). We therefore measured α18 mAb fluorescence intensity normalised for total junctional α-catenin levels (α18/pan-α-catenin ratio) as a proxy for molecular-level tension applied to the cadherin-catenin complex. Supporting our recoil measurements, α18/pan-α-catenin ratios increased at AJ within 5 min of applying hypo-osmotic media, and this was significantly reduced in Cav-1 KD monolayers (Fig. 2b-c). Specificity for Cav-1 was confirmed as the mechanical response was restored by expressing an RNAi-resistant Cav-1 transgene in the KD monolayers. Together, these data indicate that hypo-osmotic stimulation increases mechanical tension at AJ in a caveola-dependent fashion. This supported the notion that the junctional cytoskeleton was being altered by mechanical stimulation of caveolae.

Changes in junctional tension can reflect contributions from multiple elements of the AJ, potentially including the membrane itself as well as the cytoskeleton that physically couples to the cadherin-catenin adhesion complex. To dissect these possible factors, we inhibited Myosin II, which is responsible for the contractile forces that the cytoskeleton can directly and indirectly apply to AJ (Ratheesh *et al*., 2012; Smutny *et al*, 2010). Monolayers were treated with the Myosin II inhibitor, para-aminoblebbistatin (blebbistatin, 25 nM), prior to hypo-osmotic shock. Blebbistatin reduced baseline junctional tension as measured by recoil assays, consistent with the dominant contribution of cellular contractility to AJ tension. Importantly, hypo-osmotic stimulation did not increase junctional tension in the presence of blebbistatin (Fig. 2d). This suggested that cellular contractility might be principally responsible for the tensional response of AJ to hypo-osmotic stimulation. This was confirmed by measuring α18/pan-α-catenin ratios, which increased when control monolayers were stimulated with hypo-osmotic media, but not in the presence of blebbistatin (Fig. 2e). Although we cannot exclude a role for a change in membrane tension, these findings imply that cytoskeletal change may be the major process responsible for the caveola-dependent increase in junctional tension that occurs upon hypo-osmotic stimulation. Furthermore, since Cav-1 KD did not affect the response of NMII to hypo-osmotic stimulation, we surmise that tension was principally increased by the change in F-actin. This is consistent with earlier evidence that regulation of F-actin in the actomyosin system can increase mechanical tension, without concomitant change in NMII (Kovacs *et al*, 2011; Reymann *et al*, 2016; Teo *et al*., 2020; Wu *et al*, 2014).

### Caveolae mechano-activation engages a PtdIns(4, 5)P_2_-FMNL2 signalling pathway

We next sought to characterize the molecular pathway that reinforced the junctional actin cytoskeleton when caveolae were stimulated with hypo-osmotic media. Recently, we reported that the chronic depletion of caveolae activated a lipid-based signalling pathway that regulated the actin cytoskeleton (Teo *et al*., 2020). Specifically, Cav-1 RNAi increased phosphoinositide-4, 5-bisphosphate (PtdIns(4, 5)P_2_) at AJ membranes, which directly recruited the FMNL2 formin to promote actin assembly at the junctional cortex. We therefore examined whether this pathway might have a physiological role when pre-existing caveolae were acutely disassembled by mechano-stimulation.

PtdIns(4, 5)P_2_ was visualised by transiently expressing a location sensor bearing the pleckstrin homology (PH) domain of phospholipase Cδ (mCherry-PH). In control monolayers, mCherry-PH localised primarily to the plasma membrane, being clearly evident at cell-cell contacts. Junctional PtdIns(4, 5)P_2_ levels increased by ∼20% within 5 min of adding hypo-osmotic media (Fig. 3a-b and Fig. S3c). This response did not occur, however, in Cav-1 KD cells, although baseline levels were higher, as previously reported (Teo *et al*., 2020). Reconstitution of KD cells with an RNAi-resistant transgene confirmed a specific effect of Cav-1 (Fig. 3a-b and Fig. S3c). This change was relatively selective for PtdIns(4, 5)P_2_, as hypo-osmotic stress did not affect the membrane levels of PtdIns(3, 4, 5)P_3_, detected by expression of a sensor bearing the PH domain of Akt (PH-Akt-GFP) (Fig. S3a-b). Similarly, PtdIns(4, 5)P_2_ was increased by monolayer stretch (∼17%, Fig. 3c-d). Together, these findings indicated that mechanoactivation of caveolae increased the identifiable levels of junctional PtdIns (4, 5)P_2_ at AJ.

**Figure 3:**
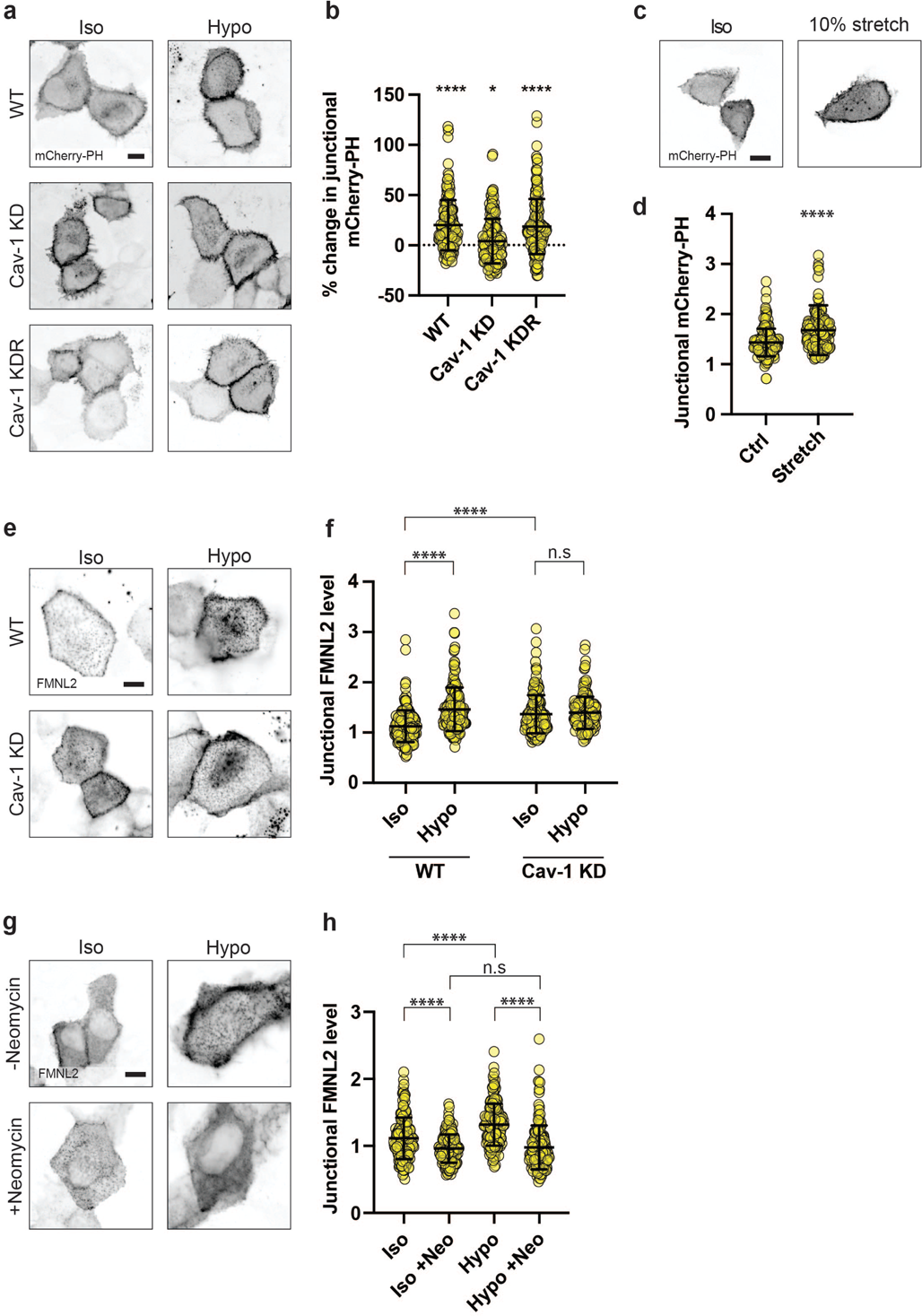
Mechanical stimulation of caveolae activates the PtdIns(4, 5)P_2_-FMNL2 signalling pathway. **(a)** Fluorescence imaging of transiently expressed mCherry-PH, a biosensor for PtdIns (4, 5)P_2_ in WT (top), Cav-1 KD (middle), and Cav-1 KDR (bottom) MCF-10A monolayers before and after hypo-osmotic shock (left and right columns, respectively) (scale bar = 10 μm). **(b)** Quantification of (a) reveals that WT monolayers are sensitive to hypo-osmotic shock, with mCherry-PH increasing by ∼20% (p = <0.0001). Cav-1 KD monolayers were relatively insensitive to hypo-osmotic shock, increasing by just ∼4% (p = 0.0823). Cav-1 KDR monolayers were resensitised to hypo-osmotic shock, with mCherry-PH increasing by ∼18% (p = <0.0001). **(c)** Fluorescence imaging of mCherry-PH expressed in control monolayers (left) and those following 5 mins of 10% stretch. **(d)** Quantification of (c) shows that mechanical stretching also increases mCherry-PH by ∼17% (p = <0.0001). **(e)** Fluorescent imaging of transiently expressed FMNL2-EGFP in WT and Cav-1 KD monolayers prior to and after hypo-osmotic shock. **(f)** Quantification of (e) shows that FMNL2-EGFP is increased by ∼30% following hypo-osmotic shock (p = <0.0001). In Cav-1 KD monolayers, baseline FMNL2-EGFP is elevated by ∼21% compared to WTs (P = <0.0001), and this is insensitive to further change with hypo-osmotic shock (p = 0.4319). **(g)** Fluorescence imaging of FMNL2-EGFP in control WT MCF-10A monolayers (top) or those pre-treated for 1 hr using 4 mM neomycin, a PtdIns(4, 5)P_2_ blocking agent (bottom). **(h)** In the absence of neomycin, FMNL2-EGFP levels were increased by ∼18% following hypo-osmotic shock (p = <0.0001). In the presence of neomycin, baseline FMNL2-EGFP levels were reduced by 14% (p = <0.0001) and were not remained unchanged by hypo-osmotic shock (p = 0.5540). All data are means ± SD. All statistical analyses calculated from N=3 independent experiments using unpaired t-tests. Points on graphs represent individual cell junctions. N.s, not significant; *p<0.05; **p<0.01, ***p<0.001; ****p<0.0001.

We then expressed FMNL2-EGFP in MCF-10A cells to assess whether it was engaged by caveola mechanoactivation. Consistent with the observed change in PtdIns(4, 5)P_2_, junctional levels of FMNL2-EGFP also increased acutely when control monolayers were stimulated with hypo-osmotic media and this response was abolished in Cav-1 KD cells (Fig. 3e-f). We then used the PtdIns(4, 5)P_2_ antagonist neomycin to test if recruitment of FMNL2 required PtdIns(4, 5)P_2_. Pre-treatment with neomycin (4 mM, 1 hr) reduced baseline levels of FMNL2 and abolished its recruitment to junctions upon hypo-osmotic stimulation (Fig. 3g-h). This supports the notion that the PtdIns(4, 5)P_2_-FMNL2 mechanism is activated when caveolae dissociate upon application of acute mechanical stress.

### Molecular stabilisation of caveolae antagonises cortical reinforcement

We next sought to test if this signalling pathway was engaged as a response to the acute disassembly of caveolae. To address this notion, we required a molecular strategy that would allow us to preserve caveolae in cells, but render them more resistant to disassembly by mechanical stimulation.

The stability of caveolae is enhanced by interactions between undecad cavin1 (UC1) repeat domains and phosphatidylserine (PtdSer) and the number of UC1 repeats influences the ability of cavin1 to stabilise caveolae (Tillu *et al*, 2018). Mammalian cavin1 contains two UC1 domains, whereas zebrafish cavin1b (*Dr*Cavin1b) has five UC1 domains, and expression of *Dr*Cavin1b in mammalian cells increased the resistance of their caveolae to disassembly upon hypo-osmotic stimulation (Tillu *et al*., 2018) (Fig. S1i). Therefore, we used lentiviral transduction to reconstitute cavin1 KD MCF-10A cells with either mouse cavin1 *(Mm*Cavin1*-*EGFP) or zebrafish cavin1 *(Dr*Cavin1b*-*EGFP), predicting that the greater number of UC1 domains would allow *Dr*Cavin1b-EGFP to stabilise caveolae against mechanical stress, compared with *Mm*Cavin1-EGFP (Fig. 4a). We then measured the change in cavin1 fluorescence at AJ before and after mechanical stimulation as an index of cavin dissociation, where negative change indicates cavin dissociation. Indeed, cavin1 dissociated from the membrane upon hypo-osmotic stress to a significantly lesser degree in *Dr*Cavin1b-EGFP cells compared with *Mm*Cavin1-EGFP cells (Fig. 4b-c and Fig. S4a). As a further control, we expressed a zebrafish cavin1 mutant construct in which four UC1 repeat domains were deleted (Δ4UC1-EGFP), bringing its UC1 complement closer to that of mouse cavin1. Stress-induced cavin dissociation in Δ4UC1-EGFP cells was similar to that in *Mm*Cavin1-EGFP cells (Fig. 4b-c and Fig. S4a). This supported the strategy that *Dr*Cavin1b, bearing a higher number of UC1 repeats, could stabilise caveolae against mechanical stress.

**Figure 4:**
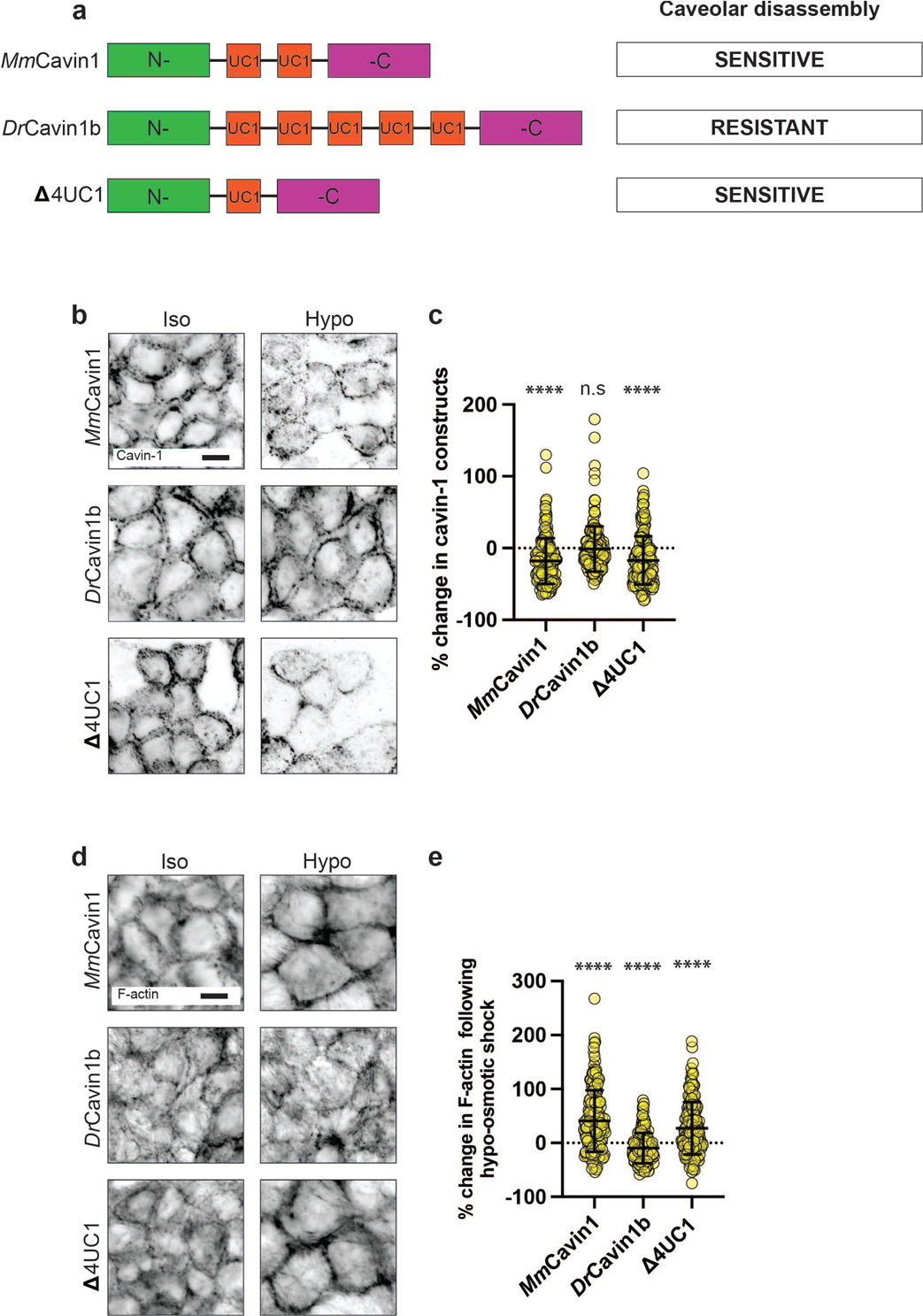
Molecular stabilization of caveolae disrupts the cytoskeletal response to mechanical stimulation. **(a)** Schematic of experimental design for molecular stabilization of caveolae through the expression of zebrafish cavin-1b *(Dr*Cavin1b) in human MCF-10A epithelial monolayers. Caveola stability is conferred through interactions between cavin-1 UC1 domains (shown in red) and phosphatidylserine (PtdSer) at the plasma membrane (PM). In contrast to the two UC1 domains of mammalian cavin-1 (top), zebrafish cavin-1b (middle) contains five UC1 domains. This confers greater stability to caveolae, resisting their disassembly following mechanical stimuli. A further cavin-1 construct containing a single UC1 domain (Δ4UC1) was adapted from zebrafish cavin-1b (bottom). This has characteristics closer to that of mammalian cavin-1 and is an additional control. **(b)** IF imaging of various cavin-1 constructs in MCF-10A monolayers which are depleted of endogenous cavin-1 by shRNA-mediated KD (scale bar = 10 μm). **(c)** Quantification of (b) showing that both mouse cavin-1 *(Mm*Cavin1; left) or a mutated zebrafish cavin-1b variant (Δ4UC1; right) (both homologous to human cavin-1) are dissociated from the PM following hypo-osmotic shock by ∼18% (p = <0.0001) and 17% (p = <0.0001), respectively. In monolayers *expressing* DrCavin1b (middle), cavin-1 is resistant to dissociation following hypo-osmotic shock, decreasing by just ∼2% (p = 0.7277). **(d)** Fluorescence imaging of F-actin in cavin-1 depleted MCF-10A monolayers expressing *Mm*Cavin1, *Dr*Cavin1b, or Δ4UC1 cavin-1 constructs (scale bar = 10 μm). **(e)** Quantification of changes to cortical F-actin levels following hypo-osmotic shock in native cavin-1-depleted MCF-10A monolayers expressing the aforementioned cavin-1 variants. In monolayers *expressing* MmCavin1 and Δ4UC1 (homologous to native cavin-1) cortical F-actin increased by ∼40% (p = <0.0001) and ∼27% (p = <0.0001), respectively. Monolayers *expressing* DrCavin1b did not show an increase in cortical F-actin (p = 0.0012). All data are means ± SD. All statistical analyses calculated from N=3 independent experiments using unpaired t-tests. Points on graphs represent individual cell junctions. N.s, not significant; *p<0.05; **p<0.01, ***p<0.001; ****p<0.0001.

We then used this molecular stabilization strategy to test how acute disassembly of caveolae contributed to reinforcing the AJ cortex against mechanical stress. For this, we reconstituted cavin1 KD monolayers with our panel of cavin1 constructs. Cavin1 KD monolayers showed mechanosensitive responses in the cortex that were altered in a fashion similar to those seen in Cav-1 KD cells. Namely, levels of junctional F-actin were increased at baseline in cavin1 KD cells, and they did not increase upon addition of hypo-osmotic media. This similarity in effect of cavin1 and Cav-1 KD supports a role for caveolae in mediating the response to hypo-osmotic stress, rather than either of these proteins independent of caveolae. Reconstitution of cavin1 KD with *Mm*Cavin1-EGFP restored their ability to reinforce the cortex, as did Δ4UC1-EGFP cells (Fig. 4d-e and Fig. S4b). Strikingly, however, cortical reinforcement was effectively abolished by the *Dr*Cavin1b-EGFP, which stabilised caveolae. This implied that the process of caveola disassembly was necessary for caveolae to reinforce the cortex in response to mechanical stress.

### Cortical reinforcement protects epithelial integrity against mechanical stress

Overall, our data indicate that the junctional cortex is enhanced by activation of PtdIns (4, 5)P_2_- FMNL2 signalling when caveolae disassemble in response to mechanical stress. In considering the potential function of this response, we hypothesized that such cortical reinforcement might increase the ability of cell-cell junctions to resist disruptive mechanical stress (Acharya *et al*., 2018; Nanavati *et al*, 2023). To test this, we stimulated cellular contractility with calyculin A, which increases mechanical tension on cell-cell adhesions, but eventually causes contacts to break.

First, we assessed if caveola disassembly was provoked by calyculin, focusing on early time points before cell-cell separation had begun. Indeed, cavin1 levels decreased upon calyculin A stimulation, as we saw with other modes of mechanical stress (Fig. 5a-b and Fig. S4c). However, this was countered when we stabilized caveolae with *Dr*Cavin1b-EGFP. Thus, after calyculin stimulation, cavin1 KD cells expressing *Dr*Cavin1b-EGFP retained greater levels of cavin1 at junctions than did KD cells reconstituted with *Mm*Cavin1-EGFP or Δ4UC1-EGFP. This indicated that our molecular stabilisation strategy operated effectively in response to contractile stress as it did for hypotonic stress.

**Figure 5:**
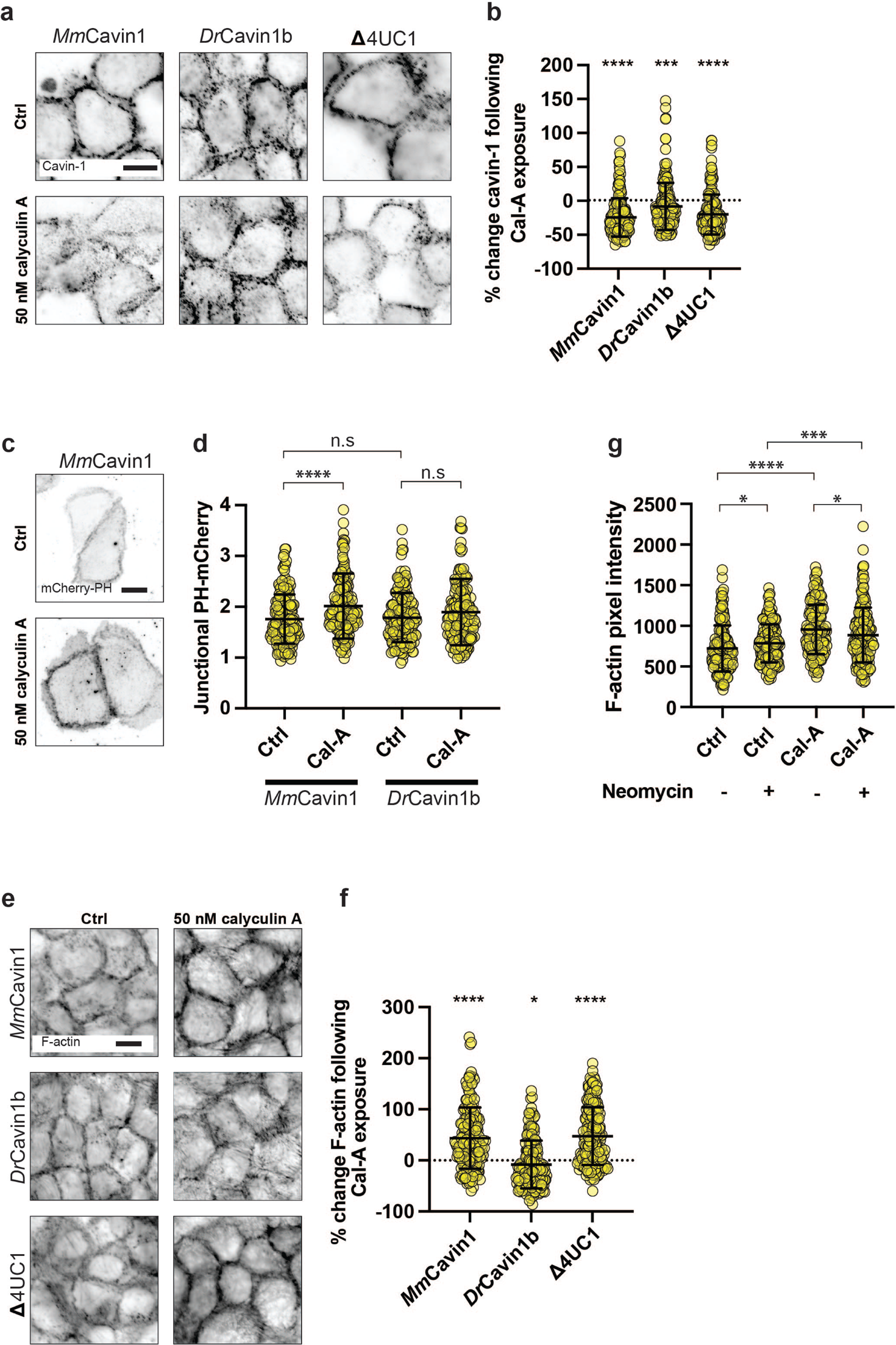
Molecular stabilization of caveolae antagonizes PtdIns(4, 5)P_2_ signalling in response to contractile stimulation. **(a)** IF imaging of various cavin-1 constructs in MCF-10A monolayers which are depleted of endogenous cavin-1 by shRNA-mediated KD (scale bar = 10 μm). **(b)** Quantification of (a) showing that both *Mm*Cavin1 and Δ4UC1 (both homologous to human cavin-1) are dissociated from the PM following 50 nM calyculin A exposure by ∼25% (p = <0.0001) and 21% (p = <0.0001), respectively. In monolayers expressing *Dr*Cavin1b, cavin-1 is resistant to dissociation, decreasing by just ∼9% (p = 0.0130). **(c)** Fluorescence imaging of PH-mCherry in cavin-1 depleted monolayers reconstituted by *Mm*Cavin1-EGFP. **(d)** Quantification of (f) shows that in cavin-1 depleted monolayers expressing *Mm*Cavin1-EGFP, PH-mCherry (PtdIns(4, 5)P_2_) is increased by ∼15% (p = <0.0001) following exposure to calyculin A. In contrast, monolayers expressing *Dr*Cavin1b-EGFP did not show an increase in PH-mCherry under the same conditions (p = 0.0734). **(e)** Fluorescence imaging of F-actin in cavin-1 depleted MCF-10A monolayers expressing the aforementioned cavin-1 variants following exposure to 50 nM calyculin A. **(f)** Quantification of (e) shows that calyculin A increased cortical F-actin by ∼44% (p = <0.0001) and 47% (p = <0.0001) *in Mm*Cavin1 and Δ4UC1 expressing monolayers, respectively. In monolayers expressing *Dr*Cavin1b, F-actin was not increased by calyculin A (p = 0.0205). **(g)** Quantification of F-actin in WT monolayers following exposure to 50 nM calyculin A. In the absence of neomycin, F-actin levels were increased by ∼32% (p = <0.0001). However, this was reduced to ∼13% in the presence of neomycin (p = 0.0009). All data are means ± SD. All statistical analyses calculated from N=3 independent experiments using unpaired t-tests. Points on graphs represent individual cell junctions. N.s, not significant; *p<0.05; **p<0.01, ***p<0.001; ****p<0.0001.

To assess downstream signalling, we used mCherry-PH to measure the PtdIns (4, 5)P_2_ response to contractile stimulation, comparing responses in cavin1 KD cells reconstituted with either Ms*Mm*Cavin1-EGFP or *Dr*Cavin1b-EGFP. Junctional PtdIns (4, 5)P_2_ increased rapidly upon calyculin stimulation in *Mm*Cavin1-EGFP cells (∼15%), but this response was compromised in *dr*Cavin1-EGFP cells (∼6%; Fig. 5c-d). This implied that contractile stress could elicit a PtdIns(4, 5)P_2_ signal at junctions as a consequence of caveola disassembly. This was accompanied by cortical reinforcement at junctions (Fig. 5e-f and Fig. S4d). Thus, junctional F-actin levels increased in response to calyculin in *Mm*Cavin1-EGFP, but not in *Dr*Cavin1b-EGFP. However, the F-actin response to calyculin was blocked by neomycin (Fig. 5g), consistent with a role for PtdIns(4, 5)P_2_ signalling. Together, these findings indicated that contractile stimulation could elicit a cortical reinforcement at junctions through caveola disassembly.

Finally, we asked if this caveola disassembly mechanism influenced the resilience of monolayers to contractile stress. For this, we followed the monolayer response to calyculin in time-lapse movies. Characteristically, monolayer integrity was lost at individual points of separation between cells, which then appeared to propagate like a fracture. Accordingly, to evaluate monolayer resilience we measured the time from addition of calyculin to when separations first appeared in the movies (Fig. 6a-b). For these experiments we used cavin1 KD cells reconstituted with either *Mm*Cavin1-EGFP or Δ4UC1-EGFP as our controls, as they showed times to initial fracture that were similar to those in wild-type MCF-10A cells. However, molecular stabilization of caveolae by expression of *Dr*Cavin1b-EGFPaccelerated the rupture process, occurring ∼20% earlier than in the controls, as it also was when PtdIns(4, 5)P_2_ signalling was blocked with neomycin (Fig 6c). This suggests that caveola disassembly increases monolayer resistance to the tensile stress of enhanced contractility.

**Figure 6:**
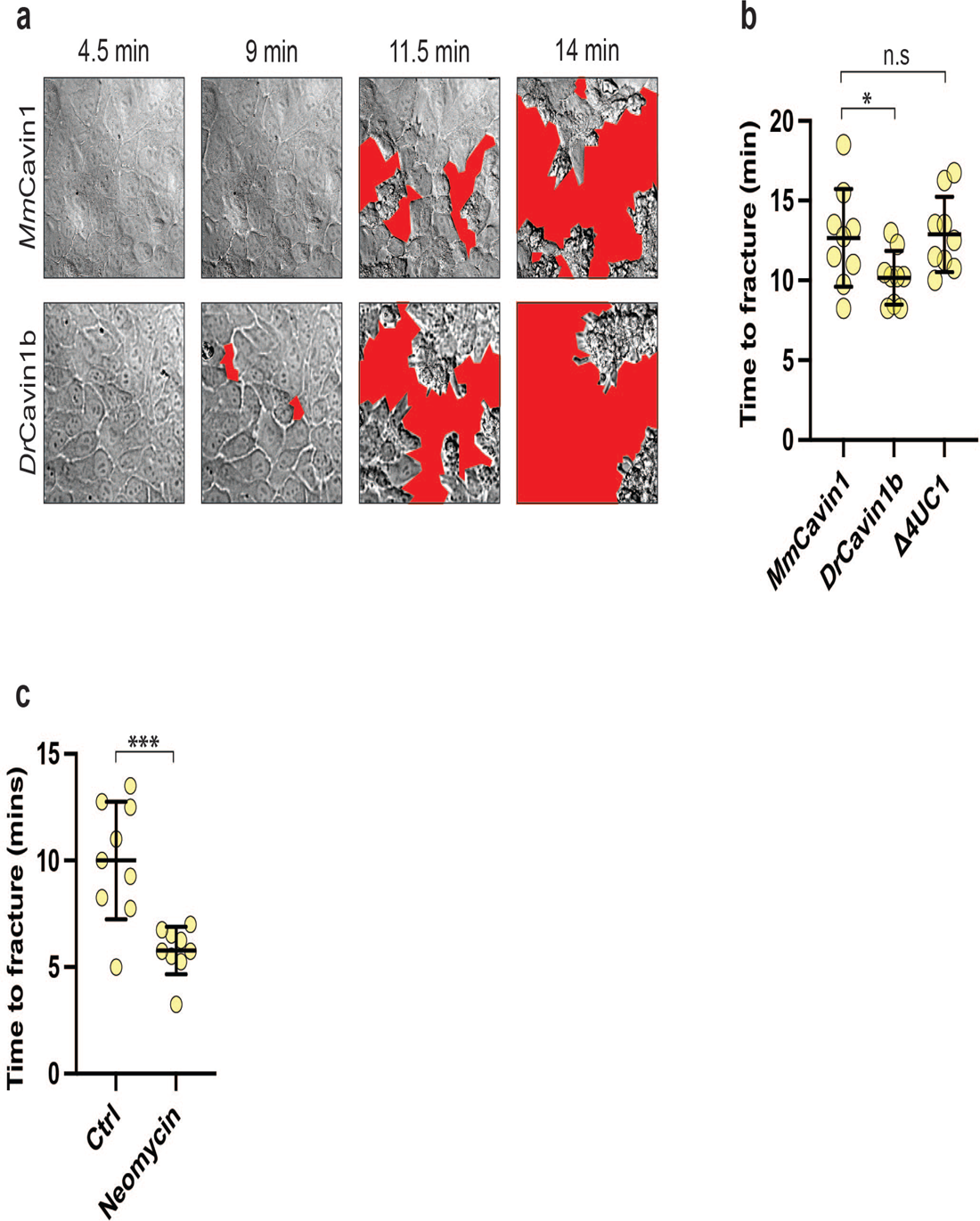
Caveola mechanotransduction confers resilience on epithelial monolayers against tensile stress. **(a)** Representative differential interference contrast (DIC) imaging of cavin-1 shRNA MCF-10A monolayers expressing *either* MmCavin1 or *Dr*Cavin1b cavin-1 variants treated with 50 nM calyculin A at 37 °C. Cell-free fractures are highlighted in red. **(b)** Quantification of (a) confirming that fracture was accelerated by molecular stabilization of caveolae with *Dr*Cavin1b. **(c)** Effect of neomycin on fracturing times in WT monolayers exposed to 50 nM calyculin A. All data are means ± SD. All statistical analyses calculated from N=3 independent experiments using unpaired t-tests. Points on graph represent individual monolayers. n.s, not significant; *p<0.05; **p<0.01; ***p<0.001; p<0.0001). Scale bar = 10 μm

## Discussion

Caveolae respond to mechanical stress by disassembly of the cavin protein coat, accompanied by partial or complete flattening of these organelles (Sinha *et al*., 2011). This process protects cells against mechanical stress by releasing reservoirs of membrane that buffer against increased membrane tension (Cheng *et al*., 2015; Del Pozo *et al*, 2021; Lee & Schmid-Schönbein, 1995; Lim *et al*., 2017; Lo *et al*., 2016; Lo *et al*, 2015; Sinha *et al*., 2011). However, caveola disassembly also has the capacity to elicit other cellular responses by increasing the availability of lipids that activate cell signalling (Teo *et al*., 2020) or, indeed, through the released cavins themselves (McMahon *et al*, 2021; McMahon *et al*, 2019; Wu *et al*, 2023). Our current experiments now identify the actin cortex as a critical target of caveola mechanotransduction. Thus, we found that the actin cortex at epithelial AJ was reinforced in a caveola-dependent fashion when monolayers were exposed to diverse forces that induced caveola disassembly. Cortical reinforcement appeared to be mediated by the recruitment of FMNL2 in response to increased accessible levels of PtdIns(4, 5)P_2_, a pathway that we earlier observed to be activated when caveolae were chronically depleted by RNAi (Teo *et al*., 2020).

Similarly, in our present studies, cortical reinforcement in response to acute mechanical stimuli was compromised by CAV-1 RNAi. However, while these observations support a role for caveolae, they did not directly test whether disassembly of already-present caveolae was the responsible mechanism. This is because the RNAi studies could not discriminate effects due to the acute disassembly from secondary effects of chronic depletion. We addressed this by developing a molecular stabilization strategy to increase the resistance of already-present caveolae to disassembly by mechanical stimulation. This was based on earlier evidence that caveola stability is influenced by the number of UC repeats in cavin1, which directly interact with phosphatidylserine in the plasma membrane (Tillu *et al*., 2018). Indeed, we found that reconstituting Cavin1 KD cells with zebrafish cavin1, which contains 5 UC1 repeats, could stabilize caveola against disassembly by mechanical stress, compared with cavin1 transgenes that had fewer UC1 repeats. Stabilization of caveolae prevented the junctional cortex from being reinforced and blocked the activation of the PtdIns(4, 5)P_2_ pathway, as measured by changes in accessible phosphoinositide. Importantly, this was accompanied by a change in the resilience of monolayers to mechanical stress. We tested this by activating cellular contractility with calyculin, which disrupts cell-cell integrity, causing monolayers to tear apart (Acharya *et al*., 2018). This fragmentation process was accelerated when we stabilized caveolae with zebrafish Cavin1, to prevent their acute disassembly. This effect was reproduced by blocking PtdIns(4, 5)P_2_ with neomycin, supporting a role for the PtdIns(4, 5)P_2_ pathway. Together, these results argue that cortical reinforcement provides a pathway for caveolar mechanotransduction to protect cell-cell integrity against disruption by mechanical stresses.

This role for cortical reinforcement in epithelial resilience is consistent with the well-characterized contribution of F-actin to cadherin adhesion. Association of classical cadherins with F-actin is essential for effective adhesion (Noordstra *et al*., 2023) and junctional cohesion is enhanced by actin assembly and stabilization (Engl *et al*, 2014; Kovacs *et al*., 2011; Lenne *et al*, 2021).

Therefore, signalling to F-actin would provide a pathway for caveola to reinforce cell-cell junctions against mechanical stresses. Of note, in our current and earlier experiments (Acharya *et al*., 2018), cell-cell adhesion appears to be a rate-limiting factor that defines monolayer resilience, as monolayer integrity is first lost when cell-cell junctions rupture. Thus, mechanical disassembly of caveolae can be understood to provide two complementary pathways for mechanoprotection. Buffering against increased membrane tension would act at the cellular level, while reinforcement of junctions would operate at the supracellular level.

Cortical reinforcement at AJ was accompanied by an increase in the mechanical line tension of the AJ themselves, consistent with earlier evidence that actomyosin-based tension can be increased by change in F-actin, as well as Myosin II (Michael *et al*., 2016; Teo *et al*., 2020; Wu *et al*., 2014). However, this result seemed paradoxical as increased junctional tension might be anticipated to increase the vulnerability of monolayers to tensile stresses. How could cortical reinforcement overcome its accompanying increase in junctional tension? Interestingly, a similar paradox was raised for tension-activated RhoA signalling, which also supports monolayer resilience to tensile stresses, despite also increasing junctional tension (Acharya *et al*., 2018). In this latter case, the paradox was resolved by modelling, which indicated that reinforcement of cortical actin increased the tensile strength of the monolayers. A similar mechanical process may explain the protective effect when caveola mechano-activation reinforces the cortex. Thus, increased line tension within AJ may be an epiphenomenon that was compensated by enhanced tensile strength.

Our results thus identify caveola mechanotransduction as one of a number of mechanisms that epithelial cells possess to detect and respond to mechanical stresses. At AJ, other mechanoprotective mechanisms include catch bonds in the adhesive ectodomain of classical cadherins as well as at their interface with cortical actin filaments (Noordstra *et al*., 2023); and tension-activated RhoA signalling based on the assembly of an E-cadherin-Myosin VI mechanosensor (Acharya *et al*., 2018; Duszyc *et al*, 2021). This raises questions about how these diverse mechanisms may be integrated. Redundancy amongst these mechanoprotective mechanisms may account for the observation that caveola-depleted animals can preserve tissue integrity (Drab *et al*, 2001; Liu *et al*, 2008). Or these mechanisms may be linked in as-yet-unknown ways. Future research will be necessary to address these issues. However, in closing we would highlight three factors that might guide such an analysis.

First, it will be important to establish the force regimes that activate these mechanosensory mechanisms. For example, catch-bonds in the E-cadherin adhesive interaction and the association of α-catenin with F-actin were engaged at forces consistent with that generated by single Myosin II motors (∼ 5 pN) (Noordstra *et al*., 2023). These therefore fit well with molecular-level mechano-protective mechanisms. Caveola mechanotransduction potentially operates at the nm scale of the organelle itself, as it involves disassembly of the cavin coat. But the force regime for this process remains to be characterized. Second, both caveola disassembly and tension-activated RhoA signalling are engaged on minutes-long time scales, suggesting that they may represent more delayed responses to mechanical stress than catch bonds. One possibility is that they represent secondary lines of response if mechanical stress persists. Finally, cadherin-based mechanotransduction mechanisms are specific for adherens junctions. Although in the present study we focused on its impact on AJ, the capacity of caveolae to signal to the actin cortex is likely to have impacts elsewhere in the cell. It will therefore be interesting to test if the pathway that we identified is used away from cell-cell junctions to allow caveolae to confer mechanoprotection to cells.

## Acknowledgements

We thank our lab colleagues for their support throughout this project, notably Ivar Noordstra for many helpful suggestions and Julia Eckert for help with the biophysical analysis. This work was supported by funds from the NHMRC (1140090 to ASY and RGP, 1136592 to ASY, APP2016410 and APP1181135 to BMC) and ARC (DP19010287 and DP220103951 to ASY). RGP was supported by an NHMRC Fellowship APP1156489 and is now an Australian Research Council (ARC) Laureate Fellow. JWB was supported by a scholarship from the UQ Research Training Programme. Additionally, this work was supported by a research training scholarship from the University of Queensland (UQRTP). Microscopy was performed at the ACRF/IMB Cancer Research Imaging Facility created with the generous support of the Australian Cancer Research Foundation.

## MATERIALS AND METHODS

### Cell culture, drugs, and transfection

MCF-10A cells were cultured at 37 °C with 5% CO_2_ in in a medium consisting of DMEM/F-12 with GlutaMAX™, supplemented to a final concentration with 5% horse serum, 20 ng mL^-1^ human epidermal growth factor (hEGF), 0.5 mg mL^-1^, 0.5 mg mL^-1^ hydrocortisone, 100 ng mL^-1^ cholera toxin, 10 μg mL^-1^ human insulin, 50 U mL^-1^ penicillin, and 50 μg mL^-1^ streptomycin. Cells were transfected using Lipofectamine 3000 (Thermo Fisher Scientific #L300015) for the expression of plasmid constructs according to the manufacturer’s guidelines. Cells were analysed 24 hours post-transfection. Where blebbistatin was used, monolayers were exposed to 25 nM para-aminoblebbistatin for 1 hour prior to mechanical stimulation/analysis. For neomycin experiments, monolayers were exposed to 4 mM neomycin for 1 hour prior to mechanical stimulation/analysis.

### Generation of stable cell lines

Stable cell lines were produced using lentiviral transduction. To generate lentiviruses, HEK-293T cells were transfected with the expression vector, PLL5.0, and lentiviral packaging constructs, pMDLg/pRRE, RSV-Rev, and pMD.G using Lipofectamine 2000 according to the manufacturer’s guidelines. Viruses were concentrated from the supernatant after 48 hours using Amicon® Ultra-15 Centrifugal Filters (Millipore UFC-9100). MCF-10A cells were transduced with the concentrated viruses, and positive cells isolated using a FACS Aria Cell Sorter (Queensland Brain Institute, University of Queensland).

### Immunostaining

MCF-10A cells were cultured to confluence and fixed with 4% (wt/vol) paraformaldehyde (PFA) in cytoskeleton stabilisation buffer (10 mM piperazine-N,N’-bis(2-ethanesulfonic acid (PIPES) at pH 6.8, 100 mM KCl, 200 mM sucrose, 2mM ethylene glycol-bis(b-aminoethyl ether)-*N,N,N’,N’*-tetraacetic acid (EGTA), 2mM MgCl2) on ice for 20 minutes. Alternatively, cells were fixed for 10 minutes in −20 °C MeOH at −20 °C. PFA-fixed cells were permeabilised for 5 minutes in 0.25% Triton X-100 in phosphate buffered saline (PBS). Cells were then blocked for 1 hour on ice using 3% bovine serum albumin (BSA) in PBS, followed by incubation with primary antibodies diluted in 3% BSA in PBS overnight at 4 °C. Cells were washed in PBS and then incubated with similarly diluted secondary antibodies for 90 minutes at room temperature prior to final washing in PBS. Coverslips were mounted onto slides using ProLong Gold containing DAPI (Cell Signalling Technologies; Cat #8961). Immunostaining for phosphorylated proteins was performed using tris buffered saline (TBS) instead of PBS.

### Junctional laser ablation

For junctional laser ablation, MCF-10A monolayers expressing E-cadherin-mCherry were used to visualise cell-cell junctions. To sever these junctions, a Zeiss 710 Inverted LSM equipped with Mai Tai two-photon capabilities was used. Ablation was performed using a 790 nm laser with 30 iterations and 22% transmission. Images were taken at 5 second intervals until the recoil of cell vertices was complete.

### Microscopy

Cells were imaged using several microscopes. Fixed samples on glass coverslips were imaged using either a Zeiss Axio 2 epifluorescence microscope using a 40x 0.95 NA Plan Neofluar or 63x 1.4 NA Plan Apochromat objective, or a Zeiss 880 inverted confocal microscope with AiryScan using a 40x 1.3 NA, or 63x 1.4 NA Plan Apochromat objective. For imaging monolayers cultured on BioFlex™ plates, imaging was performed using a Zeiss 710 upright confocal microscope using a 40x 1.0 NA water dipping N-Achroplan objective. For TFM, imaging was performed using a Zeiss AxioObserver microscope using a 40x 0.75 NA EC-Plan Neofluar objective.

### Immunoblotting

Immunoblotting was performed using sodium dodecyl sulphate-polyacrylamide gel electrophoresis (SDS-PAGE). Protein was harvested by lysing confluent monolayers in a solution containing 0.1% bromophenol blue, 2% sodium dodecyl sulphate (SDS), 50 mM Tris-Cl (pH 6.8), 10% glycerol, 200 mM dithiothreitol (DTT), and PhosSTOP (Sigma PHOSS-RO). Note that PhosSTOP was only used for detecting non-phosphorylated proteins. Cells were scraped from the dish and transferred to 1.5 mL tubes and heated to 100 °C for 10 minutes. Proteins were loaded into 1.5 mm thick homemade polyacrylamide gels consisting of a lower running gel containing between 8% and 15% acrylamide and an upper stacking gel consisting of 4% acrylamide. Protein separation was achieved by electrophoresis at 180V (400 mA) in a running buffer consisting of 25 mM Tris Base, 192 mM glycine, and 3.47 mM sodium dodecyl sulphate. Proteins were transferred to nitrocellulose filters and blocked using either 3% skim milk powder in TBS or 3% BSA in TBS. Primary antibodies were diluted in blocking buffer and incubated with the nitrocellulose filters overnight at 4 °C. Filters were then washed in TBS containing 0.1% Tween® (TBS-T) and the secondary (horseradish peroxidase, HRP) antibodies, similarly diluted, were incubated for 90 minutes at room temperature. For visualisation, membranes were exposed to a solution which allows the HRP-conjugated antibodies to emit chemiluminescence (Thermo ScientificTM 34578).

### Application of mechanical tension in monolayers

Hypo-osmotic shock was used to induce rapid cell swelling. This consisted of regular growth media, diluted in a 1:1 ratio with dH2O, giving a concentration of approximately 150 mOsm L^-1^. Cells were exposed to hypo-osmotic medium for 5 minutes (unless otherwise stated) under usual culture conditions. If applicable, cells were recovered for 1 hour in usual growth medium under usual culture conditions following hypo-osmotic shock. Mechanical stretching was performed using the Flexcell® FX-5000TM Tension System. Cells were cultured on BioFlex® plates which have a silicone base coated in type I collagen. Unless otherwise stated, stretching consisted of cyclic flexing at a frequency of 1 Hz, to a strain of 10%, for 5 minutes. Stretching was performed at room temperature. To stimulate hypercontraction of actomyosin, growth medium was supplemented with calyculin A to a concentration of 50 nM for 5 minutes under regular growth conditions (unless otherwise stated).

### Knockdown of human cavin1 and expression of *Mm*Cavin1_EGFP, *Dr*Cavin1b_EGFP, or Δ4UC1_EGFP

Endogenous cavin1 was KD from MCF-10A cells via lentivirus-mediated RNAi using the following shRNA sequence 5’-CACCTTCCACGTCAAGAAGATCCGCGAGG-3’ inserted downstream of a U6 promoter within the PLL5.0_EGFP expression vector. This shRNA targets a translated region within human cavin1. Cavin1 variants (*Mm*Cavin1, *Dr*Cavin1b, or Δ4UC1) were inserted upstream of an EGFP sequence, with expression driven by an upstream LTR viral promoter. Plasmids were synthesised by Addgene.

### Image analysis

Image analysis was performed using FIJI (ImageJ) for fluorescent microscopy, including MATLAB for TFM microscopy. For the analysis of F-actin, a line of standardised length was drawn perpendicular to the cell-cell junction and the fluorescence of all pixels along this line quantified using FIJI. The minimum intensity of this line was subtracted from the maximum intensity as background using Microsoft Excel. For all other junctional/cortical proteins, a freehand line was drawn over the entire junction, and a duplicate of this line transferred to the underlying cytosol. A ratio of the junctional to cytoplasmic ratio was calculated to determine the level of these proteins. For junctional laser ablation, cell-cell junctions were severed, and the length of the junction measured at 5 second intervals for 60 seconds, normalised to the length of the junction at time 0. For monolayer fracturing, monolayers were imaged as a time sequence at 30 second intervals following exposure to 50 nM calyculin A. Monolayer rupture was quantified according to the first frame in which cells appeared to pull apart. Prism (GraphPad) was used to generate graphs and perform statistical analysis.

## Supplementary Figure Legends

**Figure S1.**
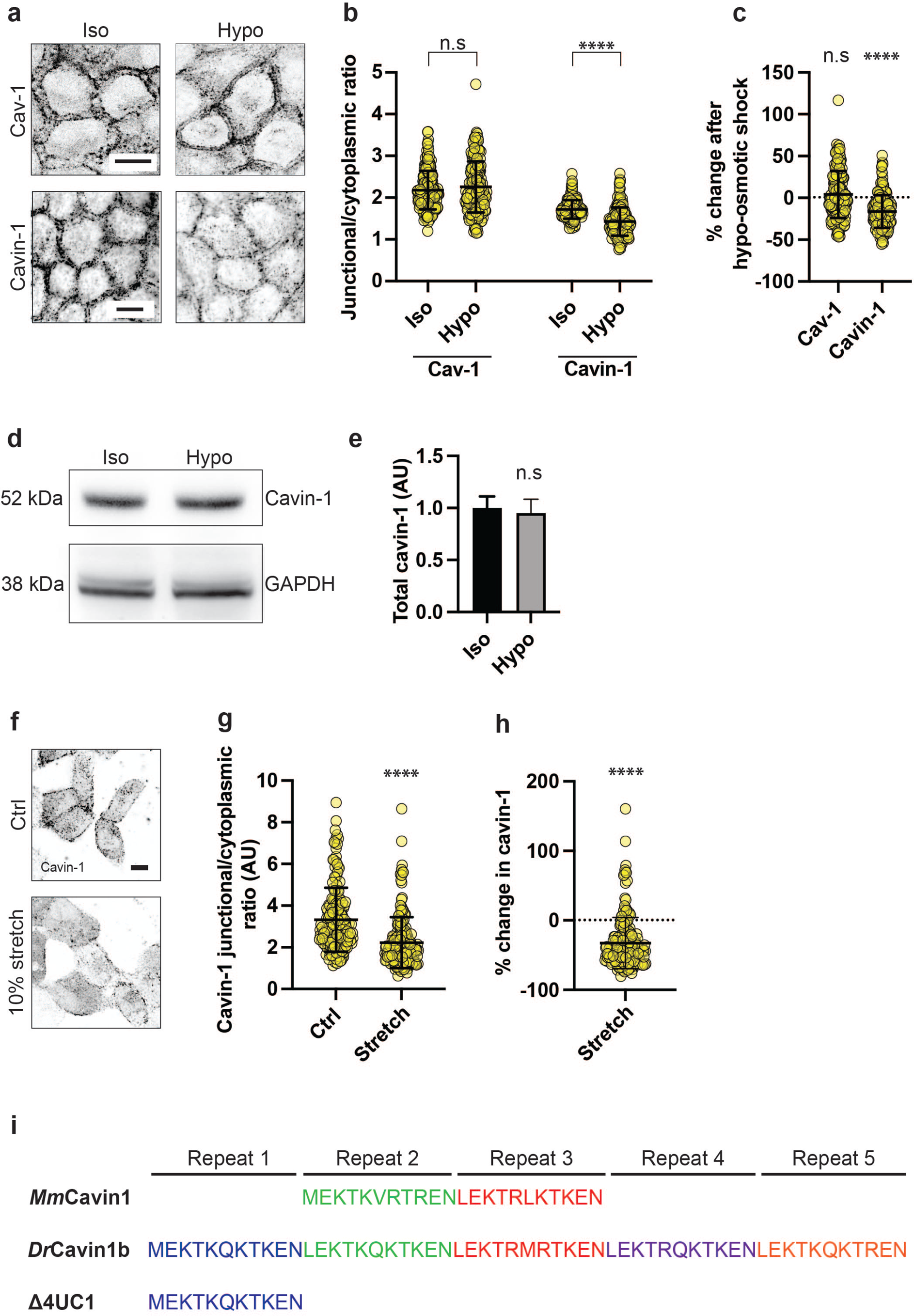
**(a)** Immunofluorescence imaging of caveolin-1 (Cav-1) and cavin-1 in iso-osmotic conditions and after 5 mins of hypo-osmotic shock (150 mOsm L^-1^). Scale bars 10 μm. **(b)** Quantification of (a) shows individual junctional to cytoplasmic ratios of these proteins. Cav-1 levels at the PM are not significantly affected by hypo-osmotic shock (p = 0.1900), whereas cavin-1 levels show a decrease of ∼17% (p = <0.0001), indicative of caveolae flattening and/or dissociation. **(c)** Quantification of (a) showing the percentage change in junctional Cav-1 and cavin-1 after hypo-osmotic shock. **(d, e)** Western Blot and analysis of cavin-1 before and after hypo-osmotic shock. Protein levels are unchanged (p = 0.2057; N = 3), indicating that cavin-1 is redistributed from the PM, rather than undergoing degradation. **(f)** Fluorescence imaging of transient cavin-1-EGFP expression in MCF-10A epithelial monolayers under control and mechanically stretched conditions (scale bar = 10 μm). **(g)** Quantification of (f) reveals that 5 m of cyclic stretching at 1 Hz to a strain of 10% promotes the dissociation of ∼33% cavin-1 from the PM (p = <0.0001), indicative of caveolar disassembly. **(h)** Quantification of (f) showing the percentage change in PM-associated cavin-1 following cyclic mechanical stretching of monolayers. **(i)** Amino acid sequences of undecad cavin1 (UC1) repeats from *Mus musculus* (*Mm*Cavin1), *Danio rerio* (*Dr*Cavin1b), and a modified *Dr*Cavin1b sequence with the deletion of four UC1 repeats (Tillu *et al*., 2018). All data are means ± SD. All statistical analyses calculated from N=3 independent experiments using unpaired t-tests. Points on graphs represent individual cell junctions. N.s, not significant; *p<0.05; **p<0.01, ***p<0.001; ****p<0.0001.

**Figure S2.**
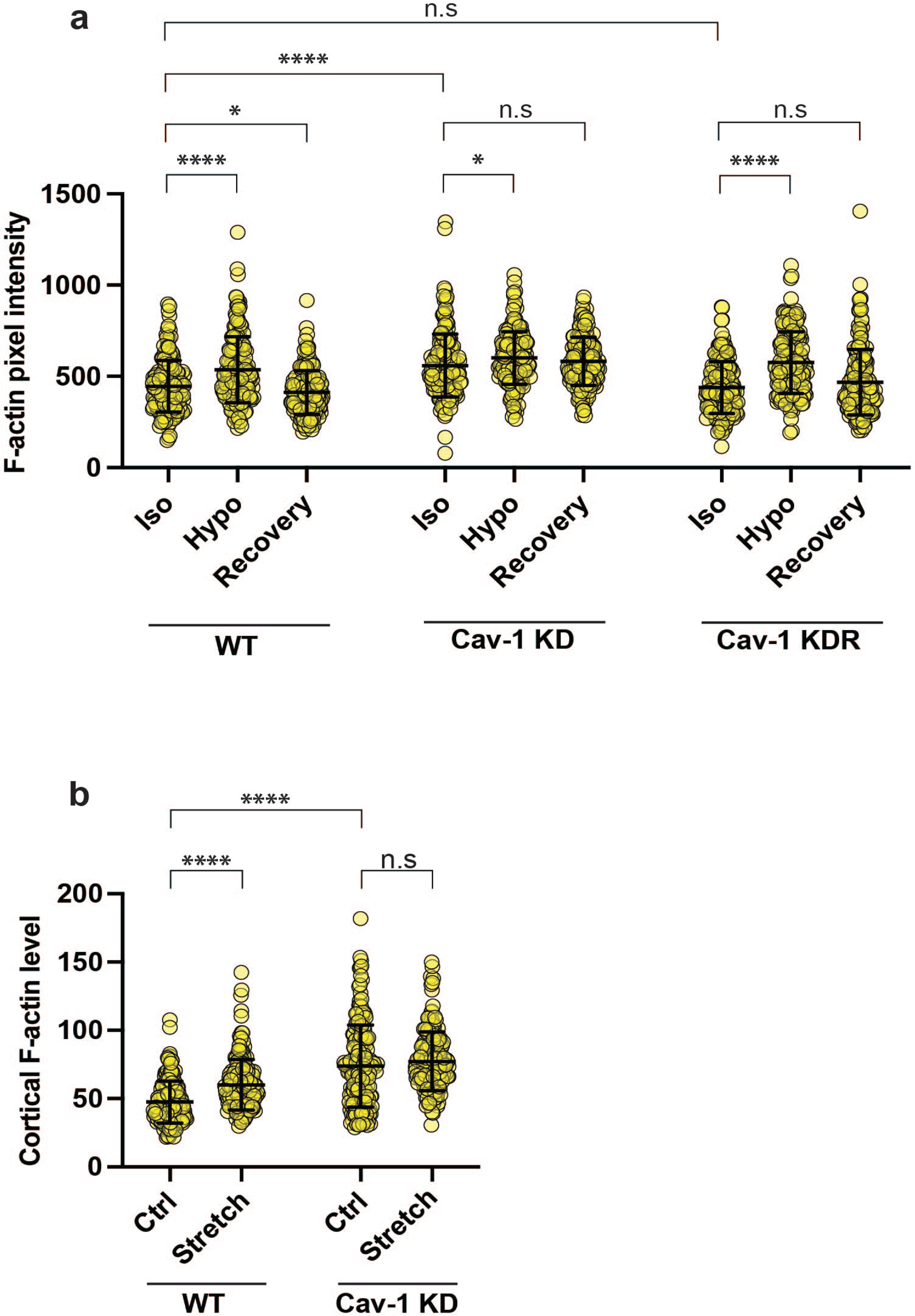
**(a)** Full dataset of F-actin changes in WT, Cav-1 KD, and Cav-1 KDR monolayers following hypo-osmotic shock (see figure 1a) showing data points from individual cell-cell junctions. **(b)** Full dataset of F-actin changes in WT and Cav-1 KD monolayers following mechanical stretch (see figure 1d) showing data points from individual cell-cell junctions. All data are means ± SD. All statistical analyses calculated from N=3 independent experiments using unpaired t-tests. Points on graphs represent individual cell junctions. N.s, not significant; *p<0.05; **p<0.01, ***p<0.001; ****p<0.0001.

**Figure S3.**
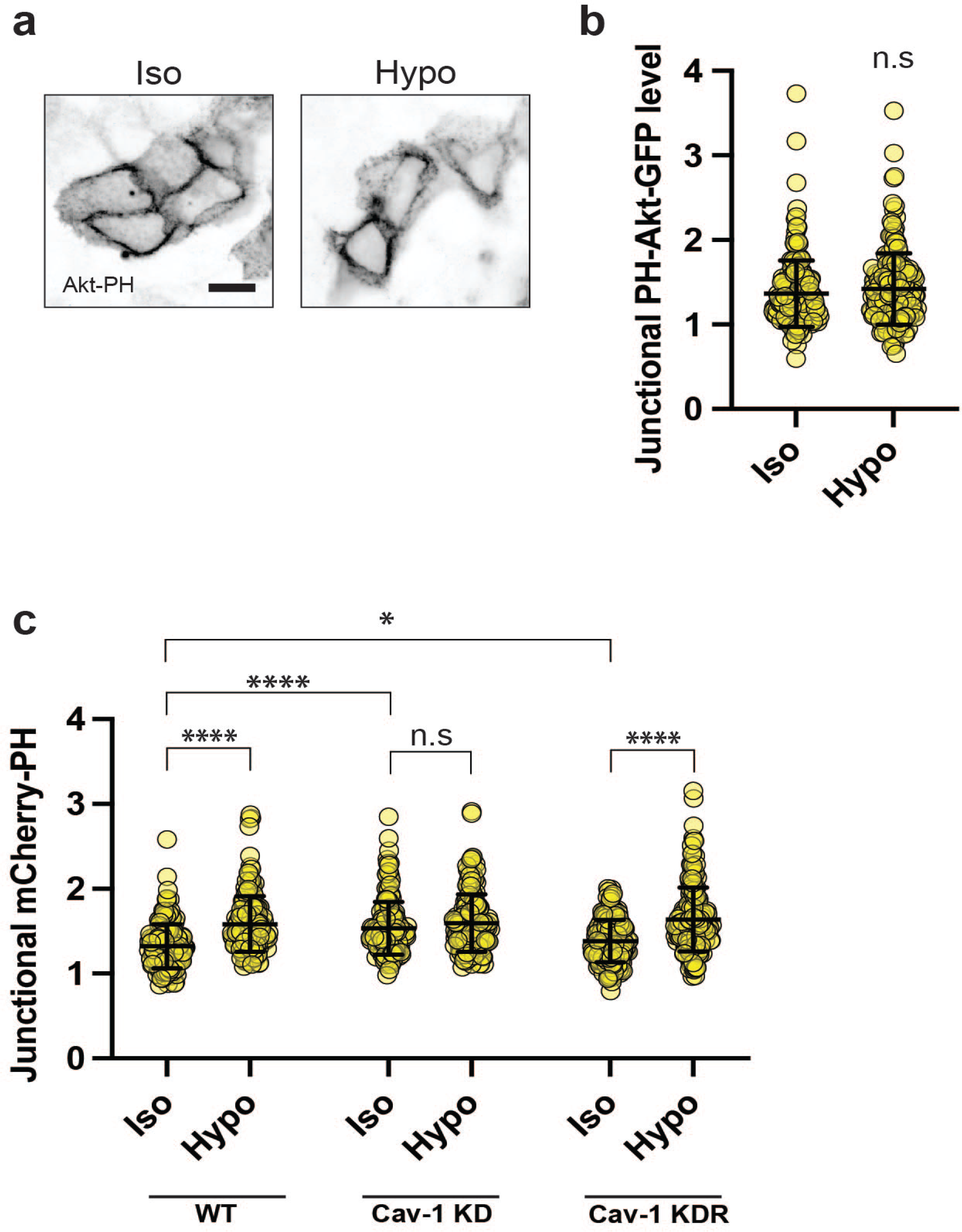
**(a)** Immunofluorescent imaging of PH-Akt-GFP, a fluorescent biosensor for PtdIns (3, 4, 5)P_3_ in WT monolayers before (left) and after (right) hypo-osmotic shock. Scale bar = 10 μm. **(b)** Quantification of (a) showing that there is no apparent change in the level of PtdIns (3, 4, 5)P_3_ (p = 0.1924), the primary precursor of PtdIns(4, 5)P2, following hypo-osmotic shock. **(c)** Full dataset of junctional mCherry-PH intensity following hypo-osmotic shock (see figure 3a) showing data points from individual cell-cell junctions in WT, Cav-1 KD, and Cav-1 KDR monolayers. All data are means ± SD. All statistical analyses calculated from N=3 independent experiments using unpaired t-tests. Points on graphs represent individual cell junctions. N.s, not significant; *p<0.05; **p<0.01, ***p<0.001; ****p<0.0001.

**Figure S4.**
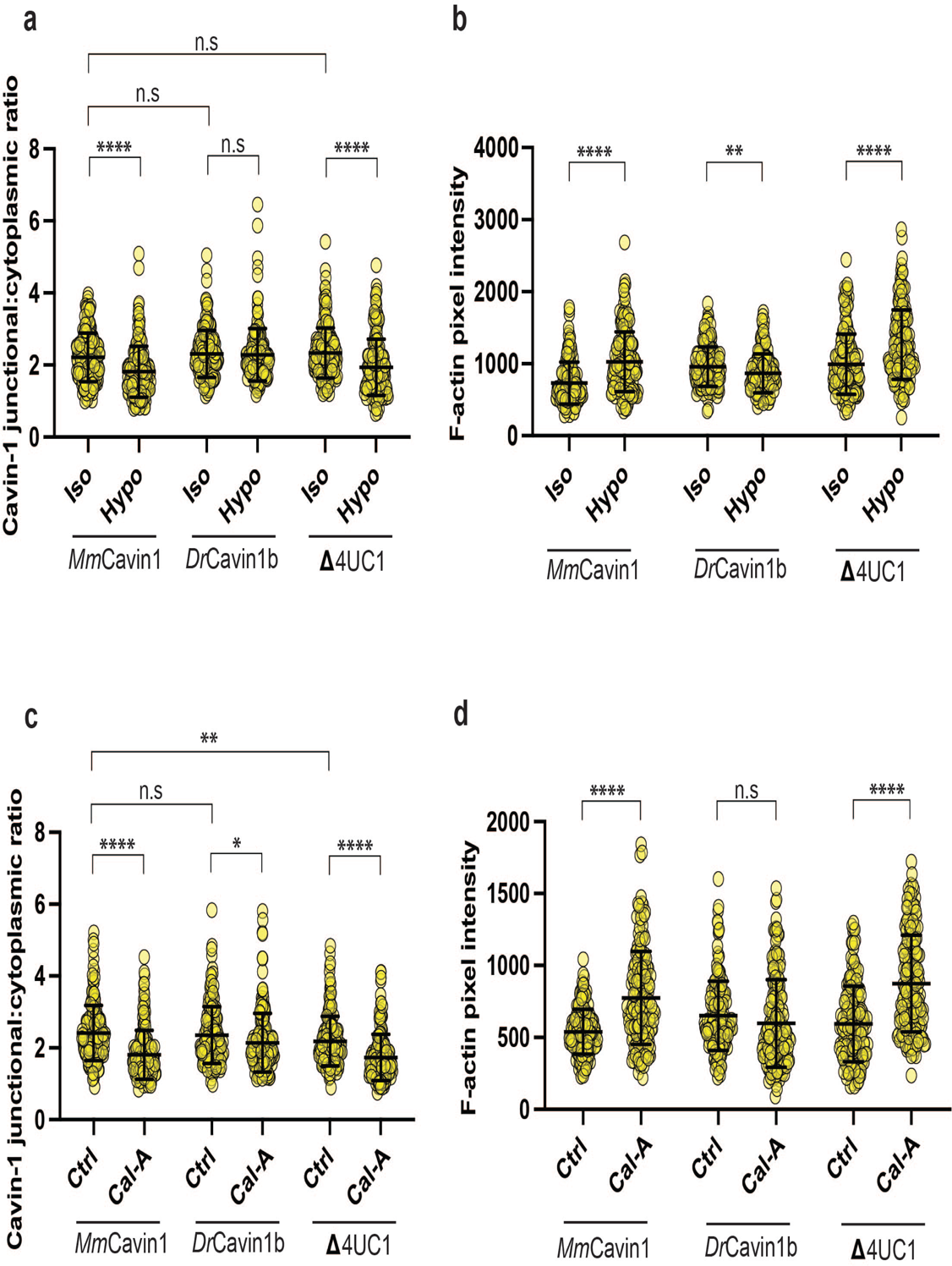
**(a)** Full dataset of cavin-1 variant dissociation following hypo-osmotic shock (see figure 4b) showing data points from individual cell-cell junctions. **(b)** Full dataset of F-actin changes in monolayers expressing either the *Mm*Cavin1, *Dr*Cavin1b, or Δ4UC1 cavin-1 variant following hypo-osmotic shock (see figure 4d) showing data points from individual cell-cell junctions. **(c)** Full dataset of cavin-1 variant dissociation following exposure to calyculin A (see figure 5a) showing data points from individual cell-cell junctions. **(d)** Full dataset of F-actin changes in monolayers expressing either the *Mm*Cavin1, *Dr*Cavin1b, or Δ4UC1 cavin-1 variant following calyculin A exposure (see figure 5e) showing data points from individual cell-cell junctions. All data are means ± SD. All statistical analyses calculated from N=3 independent experiments using unpaired t-tests. Points on graph represent individual cell junctions. n.s, not significant; *p<0.05; **p<0.01; ***p<0.001; p<0.0001).

